# Single cell migration along and against confined haptotactic gradients

**DOI:** 10.1101/2024.12.02.626413

**Authors:** Isabela Corina Fortunato, David B. Brückner, Steffen Grosser, Leone Rossetti, Miquel Bosch-Padrós, Jonel Trebicka, Pere Roca-Cusachs, Raimon Sunyer, Edouard Hannezo, Xavier Trepat

## Abstract

Directed cell migration along gradients of extracellular matrix (ECM) density –a process called haptotaxis– plays a central role in morphogenesis, the immune response, and cancer invasion. It is commonly assumed that cells respond to these gradients by migrating directionally towards the regions of highest ligand density. In contrast with this view, here we show that cells exposed to micropatterned fibronectin gradients exhibit a complex repertoire of trajectories, including directed haptotactic migration up the gradient but also linear oscillations and circles with extended periods of migration down the gradient. To explain this behavior, we developed a biophysical model of haptotactic cell migration based on a coarse-grained molecular clutch model coupled to persistent stochastic polarity dynamics. While initial haptotactic migration is explained by the differential friction at the front and back of the cell, the observed complex trajectories over longer time scales arise from the interplay between differential friction, persistence, and physical confinement. Overall, our study reveals that confinement and persistence modulate the ability of cells to sense and respond to haptotactic cues and provides a framework to understand how cells navigate complex environments.

## Introduction

The ability of single cells to follow gradients in protein concentration has been implicated in a broad range of processes in health and disease^1–4^. This type of directed cell migration is typically attributed to chemotaxis, where cells follow gradients in the concentration of soluble factors dissolved into their surrounding fluid^5,6^. However, cells can also follow gradients in the density of immobilized proteins, such as those comprising the extracellular matrix (ECM), in a process called haptotaxis. Haptotaxis was first reported by S.B. Carter in 1965, who found that fibroblasts migrate towards regions of higher substrate adhesion^7,8^. Since then, haptotaxis has been reproduced *in vitro*^9–15^ and observed *in vivo* during the immune response and cancer invasion^16–18^.

For a single cell to undergo directed migration in response to an ECM gradient, it must first detect differences in ECM density across its body, then polarize in the direction of the gradient, and finally migrate along that direction^19^. Upon contacting the ECM, integrins bind to the actin cytoskeleton through adaptor proteins such as talin, forming dynamic structures known as molecular clutches^20–22^. Actin is pulled retrogradely towards the cell center by myosin motors, exerting a centripetal force on the clutches while simultaneously polymerizing against the cell membrane to drive protrusion at the leading edge^23,24^. The dynamics of molecular clutches have been extensively studied in contexts such as cell migration^25^, adhesion^26^, and mechanotransduction^27^. However, their role in sensing ECM density gradients and driving haptotactic migration remains largely unexplored.

Besides gradient sensing, haptotactic migration is determined by the persistence of cell polarity^28–30^. If polarity is short-lived, cells are expected to respond quickly to spatial variations in the ECM density and adapt their trajectories accordingly. Conversely, if polarity is persistent, cells should sustain directed trajectories regardless of the local ECM cue, displaying an effective memory of previous polarization events. How the combination of ECM gradient sensing and polarity persistence defines cell migration trajectories is not understood. This limitation arises, in part, from the technical difficulties of generating robust ECM gradients while simultaneously measuring the key mechanical variables that govern cell migration^31–34^. In addition, despite previous theoretical approaches to ECM gradient sensing^35,36^, we lack a theoretical model that integrates such sensing with the stochastic dynamics of cell polarity.

Here we combine experimental and theoretical approaches to study the mechanics of single cells in ECM density gradients. We fabricated patterns of increasing and decreasing fibronectin density with different levels of lateral confinement. Upon adhering to the substrates, cells moved towards the regions of highest fibronectin density, demonstrating robust haptotaxis. However, after reaching the peak of fibronectin density, cells displayed a repertoire of complex trajectories depending on the level of confinement. By developing a coarse-grained clutch model, we show that this behavior can be understood in terms of the integration of cell adhesion and persistent stochastic polarity dynamics.

## Results

We developed a minimal haptotactic migration assay using soft PDMS substrates with a Young’s modulus of 12.6 kPa photopatterned with fibronectin lanes (Extended Data Fig. 1a). In a first set of experiments, the lanes were 500 µm in length and 20 µm in width, constraining the cells to migrate in 1D (Fig. 1a). The fibronectin density varied along the lane, reaching a maximum at the center, and decreasing symmetrically with an approximately exponential profile towards both ends (Fig. 1a; Extended Data Fig. 1b). We seeded single human breast epithelial cells (MCF10A) and tracked their movement for 12-20 hours immediately after seeding. We used H2B-GFP to label and track the nuclei and segmented the phase contrast image to determine the position of the leading and trailing edges. To quantify the density of each pattern, we fluorescently labeled fibronectin (Extended Data Fig. 1b; see methods). This approach allowed us to correlate the fibronectin density experienced by each cell at different time points with its velocity, length, actin dynamics and traction force.

**Figure 1.**
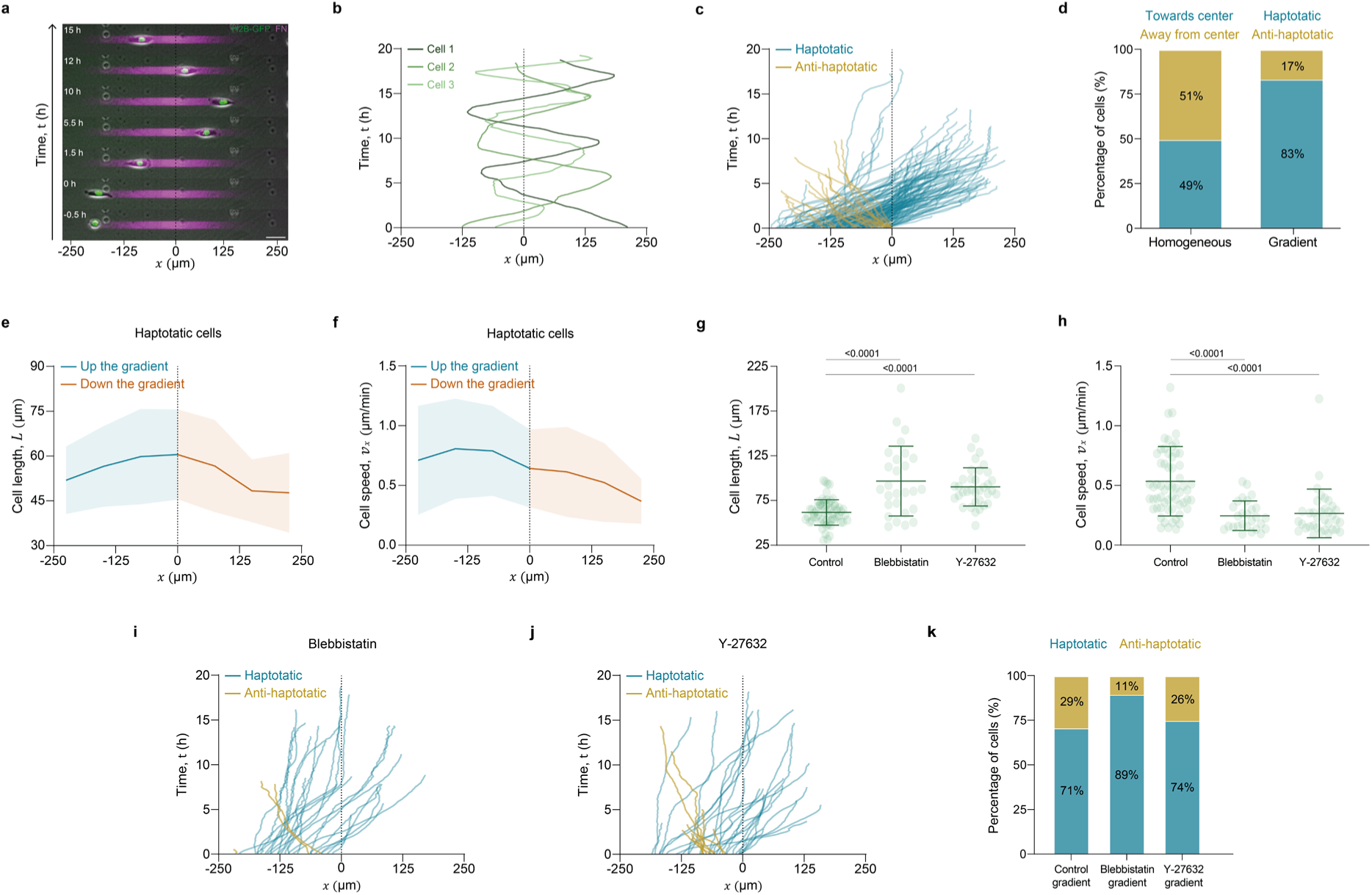
– Confined cells migrate up and down fibronectin gradients. **a**, Representative time lapse of one cell migrating on a 1D gradient. Scale bar is 50 µm. **b,** Representative trajectories of three cells migrating on 1D gradients. **c,** First run trajectories of cells migrating on 1D gradients (N=173). Color coding represents haptotactic (blue) and anti-haptotactic (yellow) cells. **d,** Percentage of cells migrating towards the center and against the center on homogeneous (N=202) and gradient (N=173) fibronectin 1D patterns. **e,** Mean length of haptotatic cells migrating on 1D gradients (N=125). Different colors label the section of migration up the gradient (blue) and down the gradient (orange). Error bars are SD. **f,** Mean velocity in the *x* direction of haptotatic cells migrating on 1D gradients (N=125). Error bars are SD. **g,** Mean length of cells migrating on 1D gradients treated with blebbistatin (N=27) and Y-27632 (N=35), compared to the control group (N=61). Error bars are SD. **h,** Mean velocity in the *x* direction of cells migrating on 1D gradients when incubated with blebbistatin (N=27) and Y-27632 (N=35), compared to the control group (N=61). Error bars are SD. For panels g-h, a significant difference between groups was identified using the Kruskal-Wallis test. To determine specific pairwise differences, Dunn’s multiple comparisons test was applied. The reported p-values correspond to Dunn’s test. **i,** First run trajectories of cells migrating on 1D gradients treated with blebbistatin (N=27). **j,** First run trajectories of cells migrating on 1D gradients treated with Y-27632 (N=35). **k,** Percentage of haptotatic and anti-haptotatic cells after incubation with blebbistatin (N=27) and Y-27632 (N=35), compared to the control group (N=61). Data without drugs were obtained from four independent experiments. Y-27632 and blebbistatin data were obtained from three independent experiments.

Upon adhering to the substrate, cells began to migrate directionally. Surprisingly, rather than simply migrating towards the region of maximum fibronectin density, they displayed oscillatory trajectories, crossing over the fibronectin peak and turning only after hours of migration down the gradient (Fig. 1a-b; Extended Data Fig. 2; Supplementary Video 1). To analyze this complex behavior, we initially focused on the first cell run, defined as the trajectory from the moment of adhesion to the first time point when cells stopped or turned. This allowed us to study the response to the gradient without the confounding contribution of the additional ECM deposited by the cells^37^. To represent cell trajectories, we took advantage of the pattern symmetry and mirrored the cellular tracks that started at *x* > 0 so that all trajectories originated from the left half of the pattern (*x* < 0, Fig. 1c). Shortly after adhering to the patterns, more than 80% of the cells polarized and migrated towards the direction of the highest protein density, indicating a robust haptotactic response (Fig. 1c-d, blue color; Supplementary Video 1). Only a minority of the cells, less than 20%, polarized against the gradient and migrated anti-haptotactically towards the region of less protein (Fig. 1c-d, yellow color). As a control, we confirmed that ∼50% of cells migrating on lanes with uniform fibronectin levels moved towards the center of the patterns (Extended Data Fig. 3; Supplementary Video 2). During this initial run, cells displayed greater elongation near the fibronectin peak (Fig. 1e). Cell velocity was highest when cells migrated up the gradient and decreased progressively down the gradient (Fig. 1f). These findings show that cells exhibit a robust initial haptotaxis, but that they can also migrate long distances against the gradient once polarized.

We next asked whether haptotaxis depends on cell contractility, similar to other modes of directed migration such as durotaxis. Previous studies have reported contrasting results. Some have suggested that haptotaxis requires the actomyosin machinery^13,14^ while others provided evidence that haptotaxis is independent of myosin and depends on lamellipodia dynamics^15^. We treated cells with blebbistatin to inhibit myosin II and with Y-27632 to inhibit Rho-Kinase. Both on lanes with homogeneous fibronectin density and on fibronectin gradients, blebbistatin and Y-27632 treatments resulted in longer and slower cells (Fig. 1g-h; Extended Data Fig. 4). However, none of these treatments diminished the initial haptotaxis response and, in the case of blebbistatin, cells tended to be more haptotactic (Fig. 1i-k). Taken together, these results show that contractility is not required for efficient haptotaxis.

We next analyzed whether the observed systematic changes in cell velocity were due to changes in the velocity of the leading edge, the trailing edge, or both. For this purpose, we examined the velocity of the leading and trailing edges of each cell and compared their values when migrating up and down the fibronectin gradient (Fig. 2a). We found that the difference in velocity at the trailing edge was non-significant, indicating that the trailing edge, on average, maintained a constant velocity regardless of the gradient sign. In contrast, the leading edge was significantly slower when migrating down the gradient. This explains the overall reduction in cell velocity (Fig. 1f) – as well as in cell elongation (Fig. 1e) – when cells migrate down the gradient.

**Figure 2.**
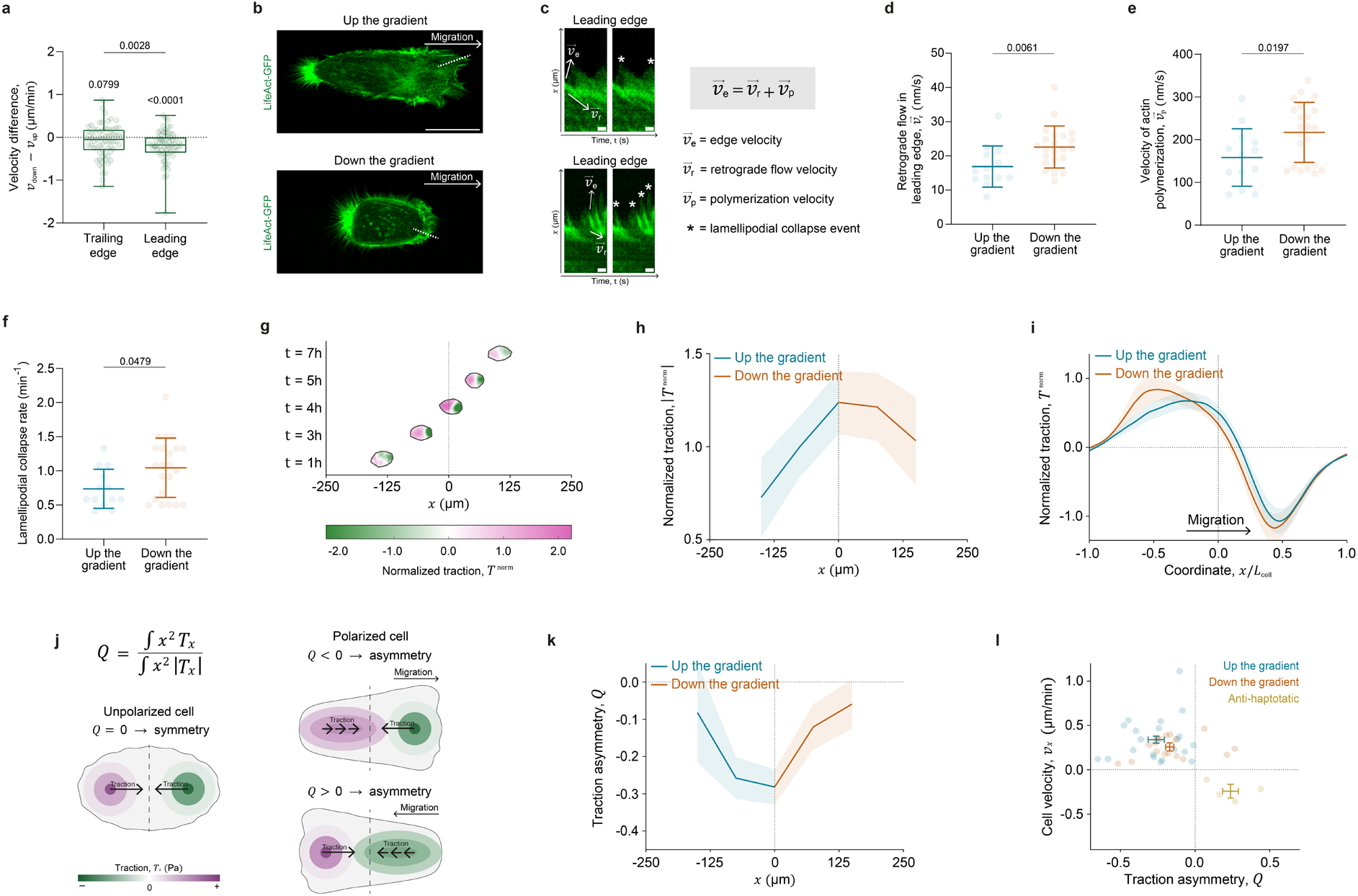
– Migration up and down the gradient involves distinct actin dynamics and traction asymmetry. **a**, Edge velocity difference of haptotatic cells migrating up and down the gradient (N=70). Error bars are SD. P-values were obtained using two types of Wilcoxon tests: one to determine if the median values in each condition are significantly different from zero, and a Wilcoxon matched-pairs test to compare the differences between paired conditions. **b,** Representative fluorescence images of MCF10A-LifeAct-GFP cells migrating up and down confined gradients. Dashed line indicates the line used to generate the kymographs in panel c. Scale bar is 20μm. **c,** Representative kymographs of leading edges in panel **b** showing the actin flow vectors and the lamellipodial collapse events. Scale bar is 1 μm. **d,** Mean retrograde flow velocity at the leading edge of cells migrating up and down 1D gradients. **e,** Mean actin polymerization velocity at the leading edge of cells migrating up and down 1D gradients. **f,** Mean rate of lamellipodial collapse events at the leading edge of cells migrating up (N=13) and down (N=22) 1D gradients. In these experiments the fibronectin fluorescence at the leading edge was similar for cells migrating up the gradient (⟨Fluorescence_Leading_ ⟩ = 164.84 ± 45.93) and down the gradient (⟨FluorescenceLeading ⟩ = 168.61 ± 31.81). Error bars are SD. P-values were obtained using Mann-Whitney test. **g,** Time lapse of the normalized traction map of a representative cell migrating up and down a gradient. **h,** Mean normalized traction for cells migrating up (N=23) and down (N=18) gradients. Error bars are SEM. **i,** Mean profiles of the *x* component of the normalized traction forces for cells migrating up (N=23) and down (N=18) the gradients. Error bars are SEM. **j,** Illustration of the traction asymmetry and its calculation through the traction quadrupole *Q*. **k,** Mean normalized traction quadrupole in cells migrating up (N=23) and down (N=18) the gradients. Error bars are SEM. **l,** Scatter plot of average velocity and normalized traction asymmetry of cells migrating up the gradient (N=23), down the gradient (N=18) and anti-haptotactically (N=4). Error bars are SEM. Actin flow data were obtained from five independent experiments. Traction data were obtained from three independent experiments.

To investigate the actin dynamics underlying the differences in the leading edge velocities, we generated MCF10A cells expressing LifeAct-GFP (Fig. 2b; Supplementary Video 3). The leading edge displayed characteristic lamellipodial dynamics, with a clear retrograde flow and frequent protrusion and retraction events. We quantified these dynamics by measuring actin retrograde flow (*v*_*r*_), edge extension velocity (*v*_*e*_), and the number of lamellipodial collapse events per unit time (Fig. 2c). We then compared these variables for cells migrating up and down the gradient. Since actin dynamics can depend both on the local fibronectin density and its gradient, we focused on two populations that experience the same fibronectin density at the leading edge, so that the only varying parameter was the sign of the gradient. We found that the actin retrograde flow at the leading edge was significantly faster in cells migrating down the gradient (Fig. 2d). Actin polymerization velocity (*v*_*p*_), which we obtained by subtracting the actin retrograde flow from the leading edge velocity, was also increased in cells migrating down the gradient (Fig. 2e). In addition, these cells displayed faster leading edge dynamics characterized by more frequent lamellipodial collapse events (Fig. 2f). Together, these data reveal that cells migrate down the gradient using an increased polymerization velocity that partially compensates the increase in retrograde flow and the more frequent lamellipodial collapses.

To further characterize the mechanics of cell migration in our system, we measured traction forces exerted by the cells on both homogeneous and gradient substrates. From the onset of migration, tractions were distributed mainly at the leading and trailing edges, forming contractile force dipoles (Fig. 2g; Extended Data Fig. 5a-b). Tractions increased in magnitude during the first 8 hours of cell migration, both in uniform and gradient substrates, indicating a transient adaptation to the substrate (Extended Data Fig. 5a). To normalize out this transient adaptation and isolate the contribution of the ECM gradient to the traction force, we divided the time evolution of the tractions exerted by each cell on gradient substrates by the average time evolution of tractions exerted by cells on uniform substrates (see methods). The resulting normalized tractions (*T^norm^*) increased with the ECM density, peaked at the center of the patterns, and decreased down the gradient, approximately following the ECM density profile (Fig. 2h).

We then examined the spatial profile of traction forces along the front-rear axis. Both for cells migrating up and down the gradient, these traction profiles were asymmetric with respect to the cell center. Specifically, the width and position of the traction peaks differed between the two halves of the cell, and the change in traction sign was shifted towards the cell’s leading edge. This asymmetry, which can be understood as a traction polarity, was more pronounced in cells migrating up the gradient (Fig. 2i). To quantify traction polarity, we computed the normalized traction quadrupole *Q*^38–40^. For symmetric (i.e. unpolarized) traction profiles the quadrupole vanishes, *Q* = 0, whereas for asymmetric profiles polarized towards the direction of growing *x*, the quadrupole is negative, *Q* < 0 (Fig. 2j). Conversely, if the traction field is polarized towards decreasing *x*, then *Q* > 0. Cells migrating up the gradient showed a negative quadrupole that decreased progressively as they approached the maximum of fibronectin density (Fig. 2k). When cells started to migrate down the gradient, the quadrupole remained negative but decreased in magnitude, becoming nearly negligible close to the edge of the patterns. We studied how the traction quadrupole is quantitatively related to cell velocity by binning all cells in three groups: cells migrating up the gradient, cells migrating down the gradient (after migrating up the gradient), and anti-haptotactic cells (i.e. migrating down the gradient from the start of the experiment, labeled yellow in Fig. 1). Nearly all cells migrating up the gradient and down the gradient displayed a positive velocity and a negative quadrupole. Hence, when plotted on a *Q* − *v* diagram, they occupied the top-left quadrant (Fig. 2l). Conversely, anti-haptotactic cells showed negative velocity and positive quadrupole, thus populating the opposite quadrant of the *Q* − *v* diagram. Overall, cells moving up the gradient tended to display faster velocity and a lower quadrupole than those migrating down the gradient. These data show that cells remain mechanically polarized as they move up and down the gradient and that their velocity is proportional to their mechanical polarity.

To explore the biophysical basis of these migratory behaviors, we developed a minimal model that integrates ECM gradient sensing with persistent cellular dynamics driven by active protrusions. Focusing on the early migration dynamics of cells that are initially unpolarized, we considered a generalization of the one-dimensional molecular clutch model with protrusion growth at both the front and back of the cell (Fig. 3a, SI Text)^30,41^. In this model, adhesive interactions with the fibronectin coating of the substrate give rise to an effective friction coupling the actin flow to the substrate. For systems with homogeneous fibronectin density, our model predicts that, at fixed actin polymerization velocity *v*_p_, the actin retrograde flow *v*_r_ decreases with increasing fibronectin density (Fig. 3a; SI text). To test this, we measured actin flows on homogeneous patterns with varying fibronectin density, and indeed found a decrease of the retrograde flow velocity with fibronectin density (Fig. 3b). Assuming that the actin polymerization has a persistence of timescale τ, our model recovers the persistent migration dynamics on homogeneous patterns (Fig. 3c; Extended Data Fig. 3c).

**Figure 3.**
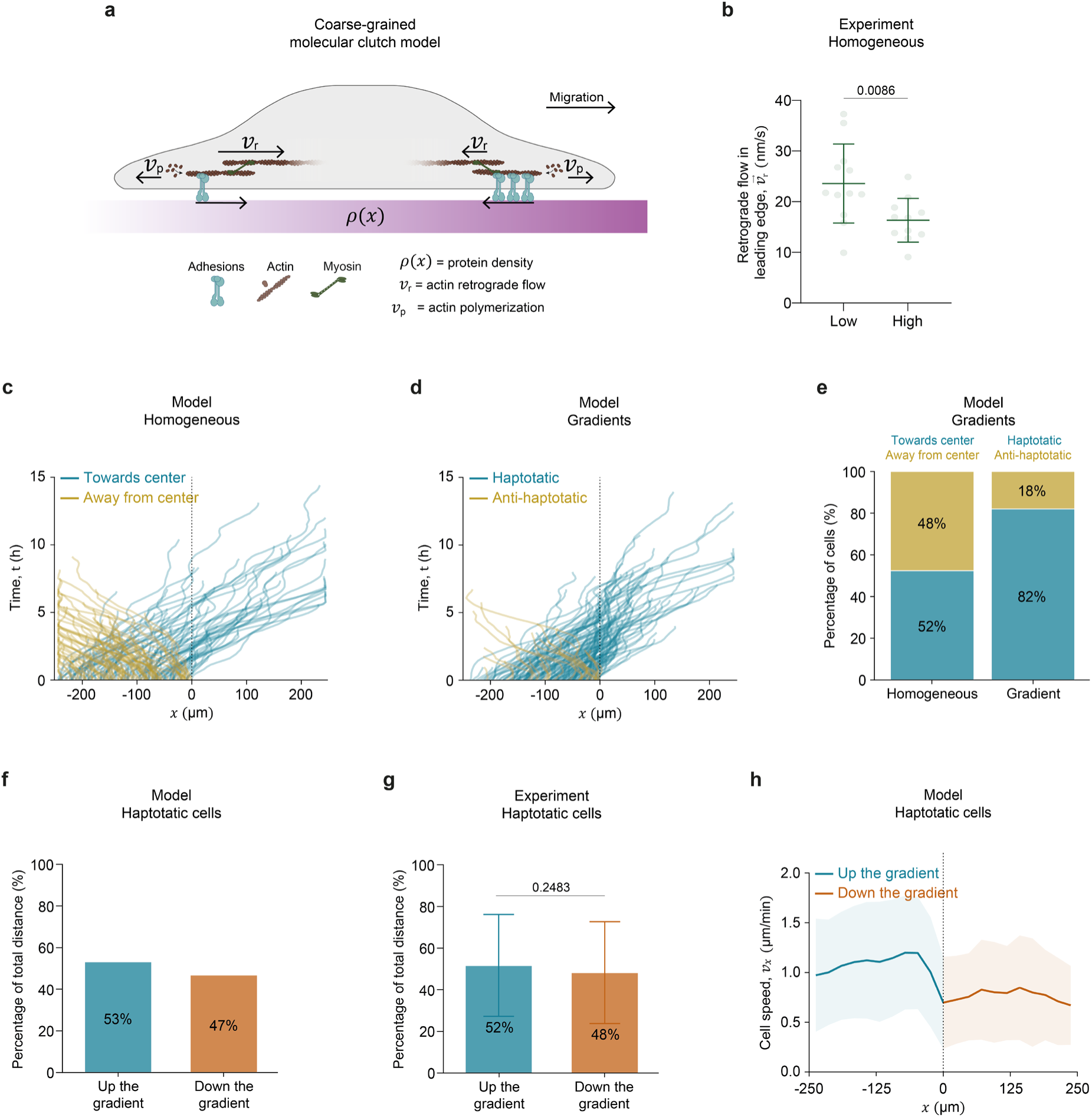
– Molecular clutch model of haptotactic cell migration. **a**, Illustration of the coarse-grained molecular clutch model. **b,** Experimental mean retrograde flow velocity at the leading edge of cells migrating on homogeneous patterns with low (N=12) and high (N=11) levels of fibronectin fluorescence. Error bars are SD. P-value obtained with Mann-Whitney test. Data were obtained from five independent experiments. **c,** Simulated first run trajectories in homogeneous fibronectin patterns with the same number of cells and duration as in experiments. **d,** Simulated first run trajectories in gradient patterns with the same number of cells and duration as in experiments. **e,** Simulated percentage of cells migrating towards the center and against the center in cells migrating on homogeneous patterns and gradients of protein density. **f,** Simulated percentage of total distance migrated by haptotatic cells when moving up and down a gradient of protein density. **g,** Experimental percentage of total distance migrated by haptotatic cells when moving up and down the gradient patterns. N=125. P-values obtained using Mann-Whitney test. Data obtained from four independent experiments. **h,** Simulated mean velocity in the *x* direction for haptotatic cells migrating on gradients of protein density. Model parameters are τ ≈ 2h, *D* ≈ 800 μm^2^h^−3^ and χ ≈ 7 × 10^9^ μm^4^/ng (SI text).

Next, we explored the consequences of differences in the concentration of fibronectin ρ between the front and back of the cell, i.e. the position-dependent gradient ∇ρ(*x*). Based on our model, this gradient gives rise to different friction between the front and back, without the need to assume any additional mechanisms, such as mechano-sensitive couplings between stress and polymerization dynamics^26,27^. This friction difference leads to higher productive forces in the protrusion further up the gradient, resulting in faster protrusion growth and net motion up the gradient. Consistent with the experiments, this friction-based mechanism does not rely on the contractility of the cell but is a purely passive mechanism. Furthermore, this friction-based mechanism predicts that traction forces should be highest in the regions of highest fibronectin density, again consistent with the experiments (Fig. 2h).

To quantitatively test our model, we focused on predicting the stochastic trajectory dynamics. To this end, we coarse-grained the model to obtain a stochastic equation of motion for the cell center *X* (SI text):

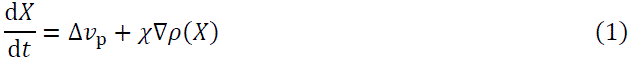

where Δ*v*_p_ is the difference in polymerization velocity between the two cell edges. We assume Δ*v*_p_ to follow persistent dynamics dΔ*v*_p_/d*t* = −Δ*v*_p_/τ + ξ(*t*), where ξ(*t*) is a Gaussian white noise with 〈ξ(*t*)〉 = 0 and 〈ξ(*t*)ξ(*t*′)〉 = 2*D*δ(*t* − *t*^′^). The coupling to the gradient is quantified by the parameter χ, which is related to the microscopic parameters of the system (SI text). Based on Eq. (1), we simulated stochastic migration trajectories on protein density gradients (Fig. 3d). Having inferred the persistence time τ ≈ 2 h and noise amplitude *D* ≈ 800 μm^2^h^−3^from trajectories of the independent experiments on uniform substrates (Extended Data Fig. 3c)^42,43^, the only remaining fit parameter is the gradient coupling χ. We find that for sufficiently large χ, the model indeed predicts trajectories strongly biased to move up the gradient and match the percentage of initially haptotactic cells to that measured in the experiment (Fig. 3d-e). Having fitted this parameter, our model makes a number of additional predictions. Firstly, once cells reach the center of the lane, due to their persistent polarity dynamics, they continue to migrate down the gradient. Quantitatively, our model then predicts that cells cumulatively migrate almost as long a distance down the gradient as up the gradient (Fig. 3f), in agreement with the experiment (Fig. 3g). Secondly, our model predicts the observed decrease in cell velocity when cells migrate down the gradient compared to their migration up the gradient (Fig. 3h), caused by the stronger frictional interaction at the trailing edge when moving down the gradient. Thirdly, our model predicts maximal traction forces and maximal cell length at the location of highest fibronectin concentration, where productive protrusion forces are the highest (SI text), consistent with the experiments (Fig. 1e; Fig. 2h). Together, these results demonstrate that a passive friction-based mechanism is sufficient to quantitatively capture the observed cellular responses to fibronectin gradients.

To further understand the mechanisms of cell migration down the gradient, we hypothesized that this behavior was related to the spatial cell confinement. Indeed, once cells become highly polarized, a 1D pattern requires a 180° flip in front-rear polarity for a cell to change its direction of migration. In contrast, a gradient in 2D could potentially allow cells to reorient their polarity more effectively through more gradual shifts in orientation. To test this idea, we progressively released the confinement by studying lanes with increasing width in the *y*-direction, orthogonal to the gradient in the *x*-direction, which remained unchanged (Fig. 4a; Supplementary Video 4). We found that the initial probability of a cell to move up the gradient is independent of the lane width, consistent with the model prediction (Extended Data Fig. 6). Interestingly, however, we experimentally observed that the long-term migration dynamics were strongly affected by the release of confinement. On 40 µm wide patterns, we observed oscillatory trajectories involving migration up and down the gradient just as in the narrowest lanes of Fig. 1a-c (corresponding to a width of 20 µm). In contrast, in quasi-2D lanes (250 µm in width), cells did not migrate down the gradient but rather made a 90° turn, migrating along the ridge defined by the maximal fibronectin density (Fig. 4b). For intermediate lane widths of 60 µm and 80 µm, cells exhibited complex patterns, including circular rotations around the midline, where fibronectin concentration is maximal (Fig. 4b).

**Figure 4.**
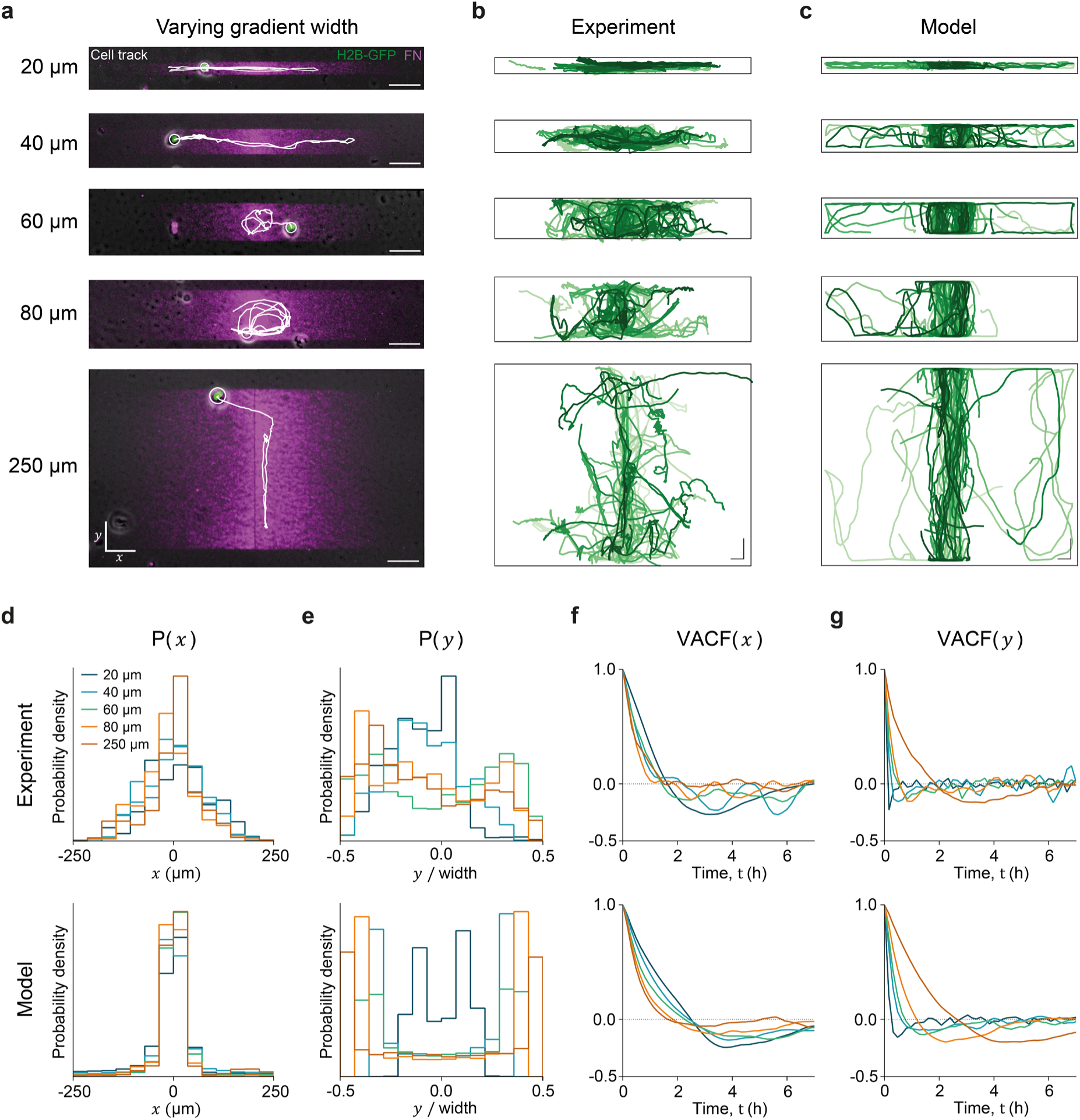
– Confinement modulates cell haptotaxis. **a**, Representative trajectories of cells migrating on gradients with widths of 20, 40, 60, 80 and 250 µm. Scale bar is 50 μm. **b,** Experimental cell trajectories monitored for 20 hours in gradients with widths of 20 µm (N=27), 40 µm (N=27), 60 µm (N=30), 80 µm (N=22) and 250 µm (N=25). Different shades of green label different individual cells. 20, 40, 60 and 80 µm data obtained from 6 replicates. 250 µm data obtained from 5 replicates. **c,** Simulated trajectories, with the same number of cells and duration as in the experiment, for each gradient width. Scale bar is 25 μm in *x* and *y*. **d,** Probability distribution P(*x*) of cell positions along the axis of the gradient (*x* axis) for different gradient widths. **e,** Probability distribution P(*y*) of cell positions in the lateral direction orthogonal to the gradient (*y* axis) for different gradient widths. **f,** Velocity autocorrelation functions in the *x* direction for different gradient widths. **g,** Velocity autocorrelation function in the *y* direction for different gradient widths. In **d-g**, top and bottom rows correspond to experiment and model, respectively.

To test whether our model of passive friction-based haptotaxis (Eq. 1) could capture these complex migration patterns, we generalized it to 2D, adapting the equation for polarity dynamics to a situation where, after initial haptotactic exploration with bi-directional protrusions, cells are polarized (with rotational diffusion, SI text). To take into account spatial 2D confinement, we consider that polarity changes are more difficult on thin lanes due to steric interactions when the width is comparable to cell size. Simulations of our model indeed capture the observed migratory patterns at the trajectory-level (Fig. 4c) and recapitulate the distributions of cellular positions in *x* and *y* (Fig. 4d-e). To further compare data and model, we examined the velocity auto-correlation functions in *x* and *y* (Fig. 4f-g). The model predicts that, with increasing width, correlations will transition from oscillatory (negative overshoot) to relaxational (decay to zero) in the *x*-direction, while the opposite trend is predicted for the *y*-direction (Fig. 4f-g). This predicted transition from oscillating in *x* to oscillating in *y* was found experimentally as shown by the auto-correlation functions computed from the experimental trajectories (Fig. 4f-g). Taken together, these results explain how the interplay of gradient sensing, polarity persistence, and physical confinement determine complex migration trajectories.

## Discussion

The study of cell migration in haptotactic gradients has traditionally emphasized that cells migrate towards the regions of highest ligand density^7–18^. Our study challenges this notion by demonstrating that cells can migrate long distances down haptotactic gradients and that the resulting trajectories depend on cell confinement and persistence. After adhering to narrow lanes of increasing and decreasing density, most cells polarized and migrated toward the highest protein density, demonstrating haptotaxis. This response was robust and independent of cell contractility. Intriguingly, after reaching the region of highest fibronectin density, most cells continued to migrate down the gradient. We captured the main features of this behavior with a coarse-grained molecular clutch model, which shows that both initial and long-term haptotactic dynamics can be explained by the interplay between friction differences at cell edges and cell persistence. Furthermore, the model predicts that cell responses to ECM gradients depend on cell confinement, which we tested by increasing the gradient width. As predicted, we found that for intermediate widths, complex trajectories such as circles emerged, while for larger widths, cells lost the ability to migrate down the gradient due to smoother polarity re-orientation compared to the effective 1D dynamics observed at narrower widths.

Cell migration down a gradient has been previously observed in chemotaxis and durotaxis^44–46^. After an initial phase of chemotactic migration along a gradient of cAMP, single Dictyostelium cells continue to migrate in the same direction when the gradient is removed or weakly inverted^44^. This effective chemotactic memory has been interpreted in terms of adaptive gradient sensing and bistability of the downstream signaling circuitry. In durotaxis, single glioma cells have also been observed to migrate against a stiffness gradient^46^. In this case, negative durotaxis is explained by a biphasic relationship between traction force and substrate stiffness and does not imply an effective memory. The mechanisms underlying migration against haptotactic gradients identified here have a different origin. We show that differences in friction between the leading and trailing edges combined with stochastic polarity persistence suffice to explain an effective memory in ECM gradients. Gradients in friction were recently proposed to drive directed cell migration in a phenomenon called frictiotaxis^47^. This type of migration differs from our observations and model in that it relies on unspecific friction rather than specific cell adhesion^47^.

While both haptotaxis and durotaxis can be explained with molecular clutch models^4,48^, mechanisms for gradient sensing are distinct. In order to sense a gradient in stiffness, cells must be able to deform the substrate, requiring the generation of a contractile force. Unlike durotaxis^48,49^, haptotaxis is not impaired by disrupting contractility (Fig. 1k). This is because haptotactic migration is driven by differences in frictional interactions between actin flows and substrate, rather than by substrate deformation. Based on our coarse-grained molecular clutch model, we found that this passive physical effect was sufficient to explain the key features of our data. In future work, our model could be extended to include active gradient sensing by coupling the cell polarity equation to local gradients as in previous work on chemotaxis^50^. Such an extended approach could help capture additional effects of our experiments such as the differences in actin flow dynamics and traction polarity for cells moving up and down the gradients (Fig. 2). However, our finding that the friction-based model already captured the initial and long-term migration on 1D lanes, as well as the complex patterns emerging when confinement is relaxed, suggests that these additional mechanisms are not dominant in our system.

Gradients in ECM density are common both in physiological and pathological conditions. Spatial variations in fibronectin density have been reported during different processes in development, such as gastrulation or limb formation^51–53^. In cancer, the aberrant accumulation of fibroblasts in the tumor microenvironment results in the secretion of large amounts of fibronectin and collagen, which vary within the tumor and decay away from it^54–56^. An accumulation of ECM is also the defining feature of fibrotic diseases and a signature of scar tissue^57,58^. Understanding how cells interpret these ECM gradients *in vivo* poses substantial challenges. One of such challenges is that an ECM accumulation not only represents a haptotactic cue but also a durotactic one. Our experimental approach allowed us to generate ECM density gradients in the absence of stiffness gradients. Using this approach, we found that haptotaxis of single MCF10a cells is far more efficient than durotaxis reported previously in the same cell type^48^. A second challenge to interpret directed cell migration in ECM gradients arises from the fact that cells *in vivo* are often confined by their 3D microenvironment^59–61^. This is well established for cancer cells during invasion or immune cells during infiltration, where migration occurs within narrow channels generated by the porous ECM, dense cellular packings, and tissue interfaces. Our technique enabled the systematic study of how confinement affects haptotactic migration. We established that confinement plays a central role in determining complex cell trajectories, including directed migration down the gradient when confinement is strong or persistent circles when it is mildly relaxed. These new results can contribute to explain the paradoxical behavior of invading cancer cells, which escape the tumor against an ECM gradient, as well as the impaired infiltration of immune cells in immuno-cold tumors^62–64^. While further work is needed to study confined haptotactic migration *in vivo*, our study *in vitro* dissects the mechanisms by which cells respond to confined ECM gradients, unveiling an unanticipated repertoire of trajectories and providing a theoretical framework that predicts their essential features.

## Methods

### Cell culture

MCF10A, MCF10A-H2B-GFP (gift from G. Charras – London Center for Nanotechnology, UK), and MCF10A-LifeAct-GFP cells were routinely maintained in filter-cap Nunc™ EasYFlask™ 25 cm^2^ or 75 cm^2^ Cell Culture Flasks at 37°C and 5% CO2 to ensure optimal growth conditions. These cells were cultured in DMEM-F12 medium (Gibco, #21331020) supplemented with 5% horse serum (ThermoFisher, #16050122), 100 U/ml penicillin-streptomycin, 20 ng/ml hEGF (Peprotech, #AF-100-15), 0.5 mg/ml hydrocortisone (Sigma-Aldrich, #H0888), 100 ng/ml cholera toxin (Sigma-Aldrich, #C8052) and 10 μg/ml insulin (ThermoFisher, #12585014).

HEK293T cells were routinely maintained in filter-cap Nunc™ EasYFlask™ 25 cm^2^ (ThermoFisher, #156367) or 75 cm^2^ Cell Culture Flasks (ThermoFisher, #156499) at 37°C and 5% CO2 to ensure optimal growth conditions. The cells were cultured in DMEM supplemented with 4.5 g/L D-glucose, L-glutamine, pyruvate (Gibco, #41966029), 10% fetal bovine serum (ThermoFisher, #10270098) and 100 U/ml penicillin-streptomycin (ThermoFisher, #10378016).

For routine passaging, cells were washed once with sterile PBS 1X (Sigma-Aldrich, #D1408) and incubated in Trypsin/EDTA (Gibco, #11590626) at 37°C and 5% CO2 for 5 minutes, in case of HEK293T, or 15-20 minutes, in case of MCF10A, MCF10A-H2B-GFP and MCF10A-LifeAct-GFP cells. When all cells were detached, complete medium was added to the T25 or T75 flask, and the cell suspension was transferred to a falcon tube. Then, cells were centrifuged for 3.5 minutes at 300 x g and further resuspended in complete medium. Cells were seeded in a new T25 or T75 previously filled with complete medium and properly diluted to keep passing them every 2-3 days.

### Stable cell line generation

MCF10A-LifeAct-GFP cells were produced by transduction of MCF10A cells with lentiviral particles generated in HEK293T after transient transfection of pLenti.PGK.LifeAct-GFP.W (Addgene, #51010) using Lipofectamine 3000 (ThermoFisher, #L3000008). Afterwards, the stable MCF10A-LifeAct-GFP cell line was obtained from a single cell colony expressing medium GFP fluorescence intensity sorted through flow cytometry (FACS) using BD FACSDiva 8.0.1 software.

### Cell seeding for experiments

For migration and traction force microscopy experiments, MCF10A-H2B-GFP cells were cultured in serum-free DMEM-F12 medium 24 hours before imaging. The serum-free medium was supplemented with 100 U/ml penicillin-streptomycin, 20 ng/ml hEGF, 0.5 mg/ml hydrocortisone, 100 ng/ml cholera toxin, and 10 µg/ml insulin, with an additional 2 mM thymidine (Sigma-Aldrich, #T1895) to inhibit cell proliferation.

When preparing the cells for the experiments, MCF10A-H2B-GFP cells were first washed once with sterile PBS 1X and then incubated for 15-20 minutes in Trypsin/EDTA at 37°C and 5% CO2. Once detached, the cells were resuspended in serum-free DMEM-F12 medium supplemented with 2 mM thymidine and transferred to a falcon tube. The cell suspension was centrifuged at 300 x g for 3.5 minutes, and the resulting pellet was resuspended in 4–6 mL of serum-free DMEM-F12 medium supplemented with 2 mM thymidine. Subsequently, 1000– 6000 cells were seeded onto 35 mm MatTek glass-bottom dishes (MatTek, #P35G-0-20-C) that had been sterilized by 10–15 minutes of UV irradiation and filled with 2 mL of serum-free DMEM-F12 medium with 2 mM thymidine.

For drug experiments, MatTek dishes were sterilized using UV light for 10-15 minutes, then filled with 2 mL of serum-free DMEM-F12 medium supplemented with 2 mM thymidine and the respective drug treatment. The drugs used were 10 µM para-Nitroblebbistatin (motorPharma, #DR-N-111) or 5 µM Y-27632 (Tocris Bioscience, #1254). As a control we used dimethyl sulfoxide (DMSO) (Sigma-Aldrich, #D8418).

For the actin flows experiments, the protocol followed was similar to that of the migration assay, with the exception that MCF10A-LifeAct-GFP cells were used. These cells were seeded in 35 mm MatTek glass-bottom dishes, which had been sterilized using UV light for 10–15 minutes. The dishes were then filled with 2 mL of serum-free DMEM-F12 medium supplemented with 2 mM thymidine and 0.02 mg rutin (Acros Organics, #132390050) to avoid GFP photobleaching.

### Preparation of soft PDMS substrates

Soft polydimethylsiloxane (PDMS) substrates were prepared following established protocols^65^. Specifically, soft PDMS (Dow, #DOWSIL™ CY 52-276) was prepared by mixing solution A and solution B in 9:10 ratio to obtain a stiffness of 12.6 kPa. The mixture was degassed for 30 minutes and kept on ice to prevent premature polymerization. Then, 150 µL of the solution was added to 35 mm MatTek glass bottom dishes pre-cleaned with air and pre-warmed at 65°C using a standard hot plate. Each dish was then spun using a spin coater (Laurell Tech., model WS-650MZ 23NPP/LITE) at 400 rpm with an acceleration of 100 rpm for 90 seconds. After spinning, substrates were kept at 65°C on a hot plate, while the remaining substrates were prepared. Once sufficient substrates were ready (between 20-40), they were incubated overnight at 65°C for complete polymerization. The substrates were then stored at room temperature and used within 1 to 1.5 months after preparation.

### Functionalization of soft PDMS substrates for photopatterning

Soft PDMS substrates were functionalized following established protocols^65,66^. Briefly, soft PDMS was incubated with 5% (3-Aminopropyl)triethoxysilane (Sigma-Aldrich, #281778) diluted in absolute ethanol for 3 minutes. After the incubation, the substrates were washed 3X with absolute ethanol, followed by 3X washes with Milli-Q® water (Milli-Q® Advantage A10 Water Purification System, Merck Milipore, #Z00Q0V0WW).

Next, the substrates were incubated for 10 minutes with a filtered bead solution containing 1:40 red fluorescent FluoSpheres™ Carboxylate-Modified Microspheres (0.2 µm, ThermoFisher, #F8810) diluted in a boric acid buffer (3.8 mg/mL sodium tetraborate [Sigma-Aldrich, #221732] and 5 mg/mL boric acid [Sigma-Aldrich, #B1934]). The substrates were then washed 3X with Milli-Q® water.

Subsequently, the substrates were incubated with 1% Poly-L-lysine (Sigma-Aldrich, #P2636) diluted in borate buffer (pH=8.3, 4.75 g/L sodium tetraborate and 3.1 g/L boric acid) for 1 hour, followed by 3X washes with 10 mM HEPES buffer (pH=8.22-8.4; Sigma-Aldrich, #H3375). Finally, the surface of the soft PDMS substrates was passivated by incubating them with 50 mg/mL solution of PEG coupled with succinimidyl valerate (Laysan Bio, #MPEG-SVA-5000) diluted in 10 mM HEPES buffer (pH=8.2-8.4) for 1 hour. The substrates were washed 3X with Milli-Q® water and stored at 4°C. Functionalized substrates were used for photopatterning within the next 2 days.

### Labeling fibronectin with fluorophore

Fibronectin from human plasma (Sigma-Aldrich, #F0895) was labeled with Alexa Fluor 647 using the Alexa Fluor™ 647 Antibody Labeling Kit (ThermoFisher, #A20186) according to the manufacturer’s instructions. The concentration of labeled fibronectin (Fibronectin-Alexa 647) was quantified using Pierce™ BCA protein assay kit (ThermoFisher, #23227) following the manufacturer’s guidelines. Spectrophotometric measurements were performed using an Infinite M200 Pro reader with i-control software. The final Fibronectin-Alexa 647 concentration was calculated as the average of 3 independent labelling procedures and their corresponding protein quantifications.

### Design of fibronectin micropatterns

Fibronectin patterns were designed using Inkscape software according to the protocols provided by Alveole (www.alveolelab.com). The designs consist of grayscale values that range from 0 to 255, with 0 (black) representing 0% protein absorption and 255 (white) representing 100% protein absorption. Homogeneous patterns were designed with a width of 20 µm and a length of 500 µm, using grayscale levels of 255, 204, 153, 102, or 51. These levels corresponded to 100%, 80%, 60%, 40%, and 20% of maximum protein absorption, respectively. Gradient patterns were designed with a width of 20 µm (1D), 40 µm, 60 µm, 80 µm or 250 µm (2D) and a length of 500 µm. Gradients featured a grayscale gradient that transitioned from a grayscale level of 0 (black) at the edges to 255 (white) at the center.

We observed that linear gradient designs produced patterns with a non-linear protein distribution, with saturation occurring near the maximum intensity. To improve the gradient profile, we measured the fibronectin density across various patterns as a function of the grayscale values used in the design. Using this calibration data, we employed a custom MATLAB script (MathWorks, USA) to determine the grayscale values required to linearize the protein gradient profile.

### Generation of fibronectin micropatterns

The soft PDMS substrates were photopatterned using the PRIMO system (Alveole, France) according to manufacturer’s guidelines. First, a photoinitiator was applied to the top of a PEG-functionalized soft PDMS substrate. The PRIMO system then projected grayscale pattern designs onto the substrate using a UV light source (λ = 375 nm). The varying UV intensities caused differential degradation of the PEG surface according to the pattern. Following irradiation, the substrates were incubated with a filtered Fibronectin-Alexa 647 solution containing 100 µg/mL fibronectin and 3.8 µg/mL Fibronectin-Alexa 647 diluted in 1X PBS for 5 minutes. The substrates were then washed three times with 1X PBS and stored at 4°C. Photopatterned substrates were used for experiments within 2 days of photopatterning.

### Fibronectin density quantification

We quantified the fibronectin density of the patterns adapting a previously established method ^33,67^. Briefly, we measured the intensity of different solutions of Fibronectin-Alexa 647 inside microfluidic PDMS channels of 2 mm length, 200 µm wide and 50 µm height. The channels were previously treated with 2% Pluronic (Sigma-Aldrich, #P2443) to prevent non-specific absorption in the internal channel walls. After perfusing the channels with different solutions of Fibronectin-Alexa 647 (0 µg/ml, 198.92 µg/ml, 397.84 µg/ml and 795.68 µg/ml), we measured the fluorescent intensity under the same conditions to those used in pattern experiments. To account for camera noise and any residual fluorescence from adsorbed proteins, we imaged the channels after rinsing them three times with PBS, and this background fluorescence was subtracted from the corresponding intensity measurements. Given that the fluorescence intensity of mercury lamps can fluctuate over time, we normalized the measurements using an auto-fluorescent homogeneous slide (Chroma, #92001) to correct for these variations. By applying the known geometry of the channels and their fibronectin concentration, we correlated the measured intensity to the surface density of fibronectin. This calibration curve was then used to convert the fluorescence intensity of the fibronectin patterns to surface density (Fig. 1b; Extended Data Fig. 1b).

### Imaging for migration and drug experiments

Before starting the time lapse, we first acquired images of an auto-fluorescent homogeneous slide to calibrate for variations in mercury lamp intensity. Then, we acquired images of the Fibronectin-Alexa 647 micropatterns. Live imaging started after setting up the microscope with thermal (37°C), CO₂ (5%), and humidity controls, and seeding the cells on the same microscope. Images were acquired every 10 minutes for 12-20 hours using a 10X 0.3 NA objective with 3 channels: phase-contrast (cell), green (nucleus) and far-red (Fibronectin-Alexa 647 micropattern). Multichannel acquisition was performed using Nikon Eclipse Ti inverted microscope with MetaMorph software.

### Imaging for actin flows experiments

Cells were imaged 4–7 hours post-seeding under controlled environmental conditions (37°C, 5% CO₂, and optimal humidity). Actin dynamics were assessed by imaging LifeAct-expressing cells at 4 second intervals over a 3 minute period using a 60X/1.40 NA objective. Additionally, the micropattern for each cell was imaged. All acquisitions were conducted using a Nikon Eclipse Ti inverted microscope equipped with a CSU-W1 confocal scanner unit.

### Imaging for TFM experiments

Similarto the migration experiments, we first acquired images of an auto-fluorescent homogeneous slide to calibrate for variations in mercury lamp intensity. Next, we captured images of the Fibronectin-Alexa 647 micropatterns and the red fluorescent beads, which were used as references for computing traction forces.

Live imaging started after setting up the microscope with thermal (37°C), CO₂ (5%), and humidity controls, and seeding the cells on the same microscope. Images were acquired every 10 minutes for 12-20 hours using a 20X 0.45 NA objective with 4 channels: phase contrast (cell), green (nucleus), red (beads) and far-red (Fibronectin-Alexa 647 micropattern). Multichannel acquisition was performed using Nikon Eclipse Ti inverted microscope with MetaMorph software.

### Cell tracking, cell segmentation and migration analysis

Cell tracking was performed using the TrackMate plugin from Fiji^68,69^ using the H2B-GFP nuclear marker expressed by the cells. Cell length was computed from the phase contrast images using a custom-made MATLAB script. Another MATLAB script integrated data on Fibronectin-Alexa 647 density, cell tracking, and cell length to create a comprehensive dataset describing cell migration throughout the live imaging period.

### Quantification of actin flows and lamellipodial collapse events

Acquired images of actin were processed with the Fiji software^69^. Kymographs were generated in 4 regions of the leading edge. In each region, actin retrograde flow (actin movement from the edges toward the cell center) and edge velocities were manually measured. The velocity of actin polymerization (*v*_*p*_) was calculated using the following equation:

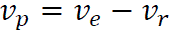

where *v*_*e*_ is the velocity of the cell’s edge and *v*_*r*_ the actin retrograde flow (Fig. 2c). To measure cell velocity, length, and fibronectin fluorescence, cell masks were created using the LifeAct-GFP signal as a reference, applying the Otsu threshold in Fiji. The masks and pattern images were oriented with the leading edge on the right side of the images and the trailing edge on the left. These masks were then used to select cells migrating faster than 0.1 µm/min and to calculate the fluorescence slope across the cell. The fluorescence slope was used to classify cells as migrating up the gradient, down the gradient or on homogeneous patterns. The lamellipodial collapse rate was manually measured by quantifying the number of actin peaks in the kymographs of the leading edge and dividing it by the total imaging time (3 minutes). The cell’s leading edge was defined as the right edge of the mask, extending 4 µm in length. Fibronectin fluorescence in the leading edge was defined by the mean fluorescence in the first 4 µm of the edge.

### Traction force data and asymmetry analysis

All traction computations and the following analyses of traction forces were carried out with custom-written MATLAB scripts. Fourier transform traction microscopy was used to measure traction forces^70–72^. The displacement fields of the fluorescence microspheres were obtained using a self-written particle imaging velocimetry (PIV) algorithm using square interrogation windows of side length 24 pixels (corresponding to about 7.8 µm) with a relative overlap of 0.75 window length. Segmented binary masks of the phase contrast images were used to extract the tractions under each cell at each time point, after dilation of the segmentation by 15 µm in all directions to include the traction profiles as complete as possible.

Traction magnitudes are the absolute values of *x*-traction, averaged over the area of a cell. Traction profiles were calculated by averaging the *x*-component of the tractions within the segmented area across the *y*-axis at every time point. The axial lengths of these profiles were normalized to unit length for each time point, and these length-normalized tractions were averaged together in groups according to the cell trajectory (haptotactic, anti-haptotactic, or homogeneous fibronectin migration) to form the averaged profiles. To remove the trend of increasing traction values with time, the traction magnitudes and profiles are further normalized. For this, the temporal traction development of cells on lanes with homogeneous fibronectin are used as reference. The magnitudes are divided by the average magnitude of the reference cells, depending on time. The profiles are divided by the maximum of the profile of the reference cells, depending on time. This creates unitless traction magnitudes and profiles which typically reach values of about 1 for the reference condition.

The normalized second moment *Q* of the 1D-traction profiles was calculated as 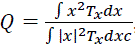 analogously to Rossetti et al.^40^, where *x* is the spatial coordinate relative to the center of mass of the traction segmentation. The moments were then averaged over the corresponding times for each cell.

### Statistical analysis

Graphs and statistical analysis were made using MATLAB, GraphPad Prism 8.0.2/10, RStudio and Microsoft Excel. Figures and figure legends describe the following statistical details: number of cells, error bars, number of independent experiments, statistical tests, and corresponding p-values. We considered the result significant when p-value < 0.05.

## Data availability

The full datasets that support the findings of this study are available from the corresponding authors.

## Code availability

All analysis procedures and codes are available from the corresponding authors.

## Acknowledgements

We thank all the members of our groups for their discussions and support. We thank A. Menéndez, S. Usieto, M. Purciolas and E. Coderch for technical assistance. We thank G. Charras (London Center for Nanotechnology, UK) for sharing cells used in this work. We thank P. Guillamat for technical advice and A. Labernardie for providing the microfluidic channels. We thank M. Gómez-González for sharing the 2D traction microscopy algorithm. Finally, we thank P. Guillamat, J. Abenza, G. Ceada, L. Faure, E. Dalaka, M. Matejčić, A. Beedle, I. Granero, O. Baguer, A. Albajar and N. Chahare for discussions. This paper was funded by the Generalitat de Catalunya (AGAUR SGR-2017-01602 to X.T.; 2021 SGR 00523 to R.S.; the CERCA Programme and ‘ICREA Academia’ awards to P.R.-C.); the Spanish Ministry for Science and Innovation MICCINN/FEDER (PID2021-128635NB-I00, MCIN/ AEI/10.13039/501100011033 and ‘ERDF-EU A way of making Europe’ to X.T.; PID2021-128674OB-I00 and CNS2022-135533 to R.S.; PID2019-110298GB-I00 to P.R.-C.); European Research Council (101097753 to P.R.-C. and Adv-883739 to X.T.); Fundació la Marató de TV3 (project 201903-30-31-32 to X.T.); European Commission (H2020-FETPROACT-01-2016-731957 to P.R.-C. and X.T.); La Caixa Foundation (LCF/PR/HR20/52400004 to P.R.-C. and X.T.); R.S. is a Serra-Hunter fellow; D.B.B. was supported by the NOMIS foundation as a NOMIS fellow, by the European Molecular Biology Organization (Postdoctoral Fellowship ALTF 343-2022) and by the Austrian Academy of Sciences through an APART-MINT Fellowship; I.C.F. acknowledges support from the European Foundation for the Study of Chronic Liver Failure; IBEC is recipient of a Severo Ochoa Award of Excellence from MINECO.

## Authors contributions

R.S., X.T. and I.C.F. conceived the project. R.S and X.T supervised the project. I.C.F. and R.S. developed the haptotatic system. I.C.F., R.S. and X.T. designed the experiments. I.C.F. performed the experiments. R.S., I.C.F., S.G., L.R. and M.B.-P. developed computational analysis tools. I.C.F. processed the experimental data. I.C.F., S.G., R.S. and D.B.B analyzed the data. D.B.B. and E.H. developed the theoretical model. D.B.B. implemented and performed the simulations. J.T. and P.R.-C. contributed technical expertise, materials, and discussion. X.T., I.C.F., R.S., D.B.B. and E.H. wrote the manuscript. All authors revised the completed manuscript.

