## Supplementary Materials for "Single cell migration along and against confined haptotactic gradients"

#### Supplementary Videos

**Supplementary Video 1 –** Representative time-lapse video of MCF10A cells migrating on confined fibronectin density gradient patterns. Cells initially move along the fibronectin gradient but continue to migrate persistently against it, resulting in oscillatory trajectories. Video rate is 15 frames per second.

**Supplementary Video 2 –** Representative time-lapse video of MCF10A-H2B-GFP cells migrating on confined homogeneous fibronectin density patterns. Cells randomly initiate migration and oscillate between the pattern extremes. The fibronectin pattern is labelled with Alexa-647. Video rate is 15 frames per second.

**Supplementary Video 3 –** Representative time-lapse video of actin flows of MCF10A-LifeAct-GFP cells migrating up (left side) and down (right side) confined fibronectin density gradient patterns. The leading edge of cells migrating down the gradient display more dynamic behaviour compared to those migrating up the gradient, with faster actin retrograde flow, increased actin polymerization, and a higher lamellipodial collapse rate. Video rate is 15 frames per second.

**Supplementary Video 4 –** Representative time-lapse videos of MCF10-H2B-GFP cells migrating on fibronectin density gradient patterns with varying widths. Upon adhesion, cells move towards the maximum protein region irrespective of the confinement level. However, the long-term migration dynamics is strongly affected by the confinement. On confined gradients (20 μm and 40 μm width) cells migrate down the gradient once they reach the fibronectin peak. For intermediate lane widths of 60 μm and 80 μm, cells exhibited non-trivial patterns, including circular rotations around the midline, where concentration is maximal. In near 2D patterns (250 μm), cells do not migrate down the gradient but rather made a 90º turn, migrating along the ridge defined by the maximal fibronectin density. Video rate is 15 frames per second.

#### Extended Data Figures

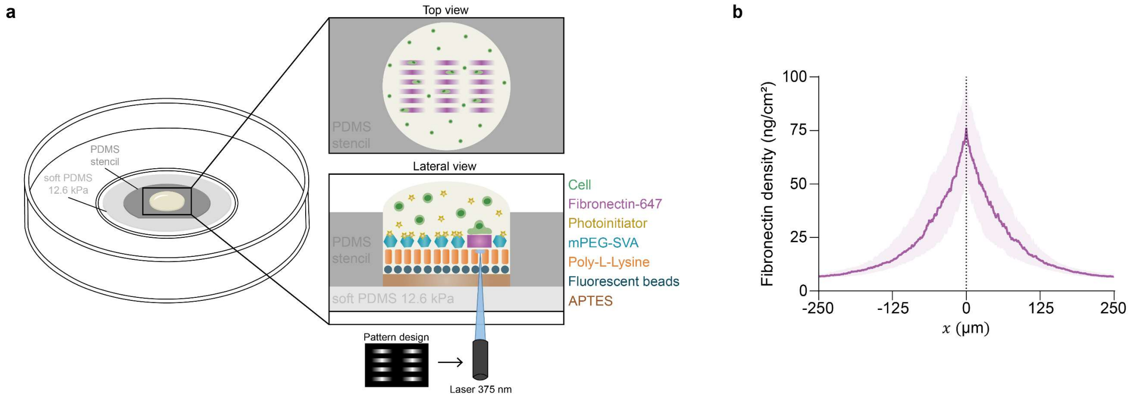


**Extended Data Figure 1 – Experimental setup. a,** Illustration of micropatterning soft elastomeric substrates by light-induced molecular adsorption of proteins (LIMAP) using PRIMO system. **b,** Median fibronectin density of 1D gradients. Error bars are 25 and 75 percentiles. N=173. Data were obtained from four independent experiments.

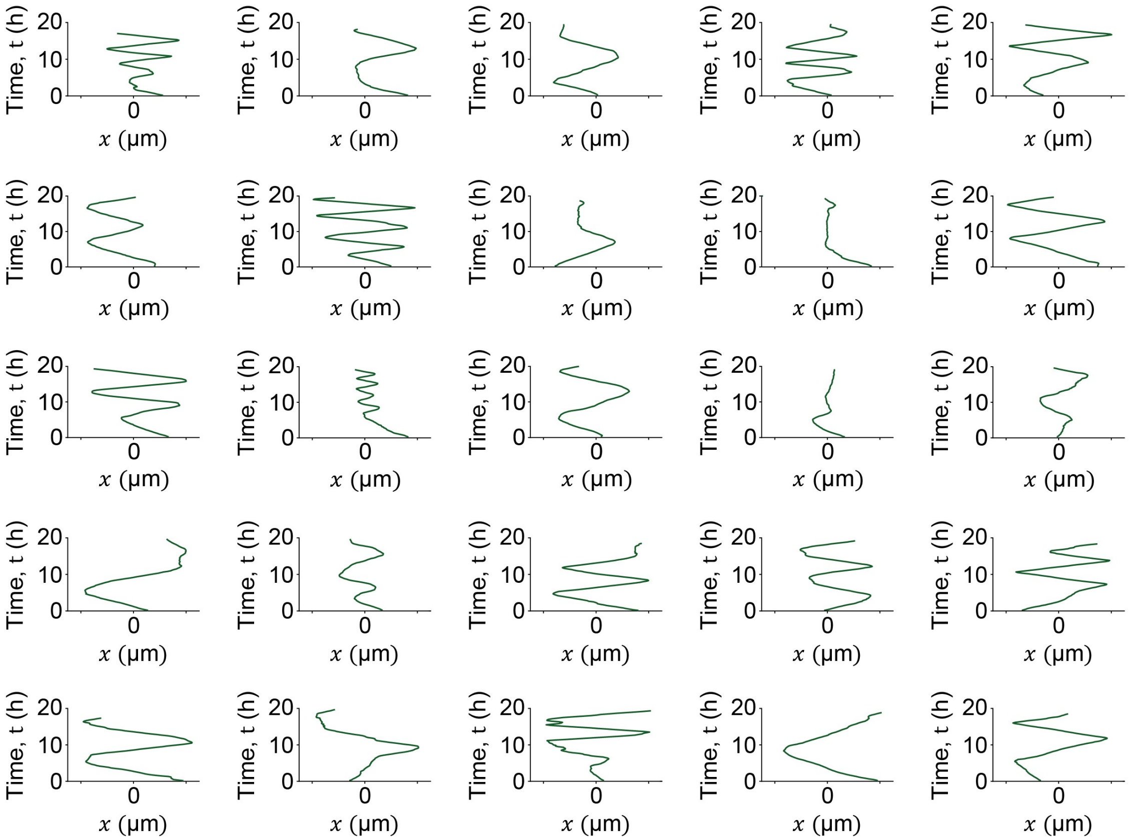


**Extended Data Figure 2 – Examples of full cell trajectories.** 25 full trajectories representative of N=173 cells migrating on 1D gradients.

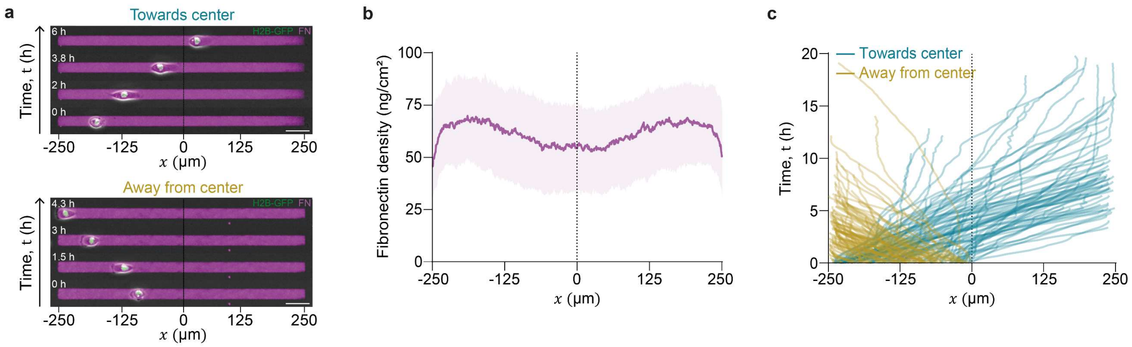


**Extended Data Figure 3 – Migration on homogenous patterns. a,** Representative fluorescence images of cells migrating on 1D patterns with homogeneous fibronectin. Scale bar is 50 µm. **b,** Median fibronectin density of 1D patterns with homogeneous fibronectin. Error bars are 25 and 75 percentiles. N=202. **c,** First run trajectories of cells migrating on 1D patterns with homogeneous fibronectin. N=202. Data were obtained from four independent experiments.

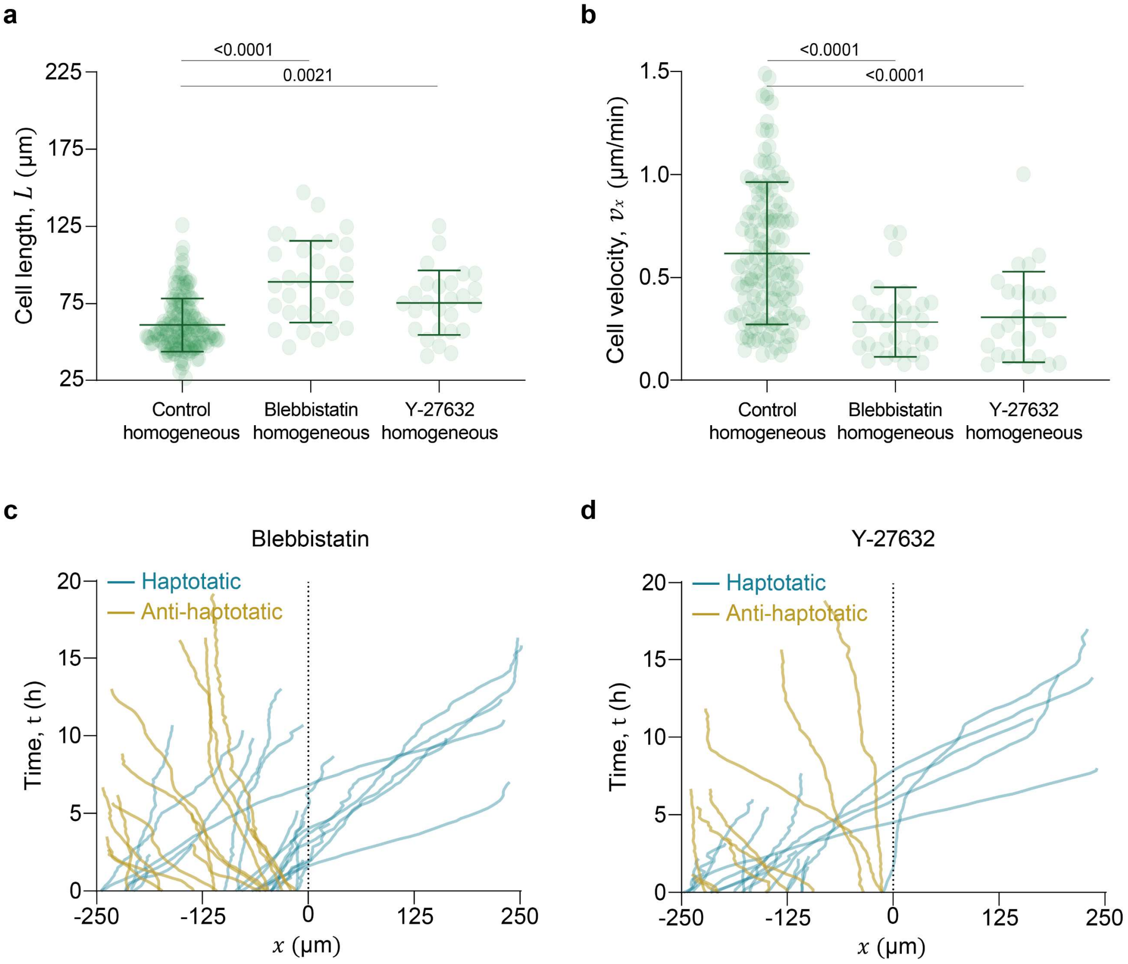


**Extended Data Figure 4 – Cell length and velocity is affected upon inhibition of myosin or ROCK in 1D patterns with homogeneous fibronectin. a,** Mean length of cells migrating on 1D patterns with homogeneous fibronectin when incubated with blebbistatin (N=32) or Y-27632 (N=26), compared to the control group (N=151). Error bars are SD. **b,** Mean velocity in the $x$-axis of cells migrating on 1D patterns with homogeneous fibronectin with treatments of blebbistatin (N=32) and Y-27632 (N=26), compared to the control group (N=151). Error bars are SD. For panels a-b, a significant difference between groups was identified using the Kruskal-Wallis test. To determine specific pairwise differences, Dunn's multiple comparisons test was applied. The reported p-values correspond to Dunn's test. **c,** First run trajectories of cells migrating on 1D patterns with homogeneous fibronectin treated with blebbistatin (N=32). **d,** First run trajectories of cells migrating on 1D patterns with homogeneous fibronectin treated with Y-27632 (N=26). Control data were obtained from four independent experiments. Y-27632 and blebbistatin data were obtained from three independent experiments.

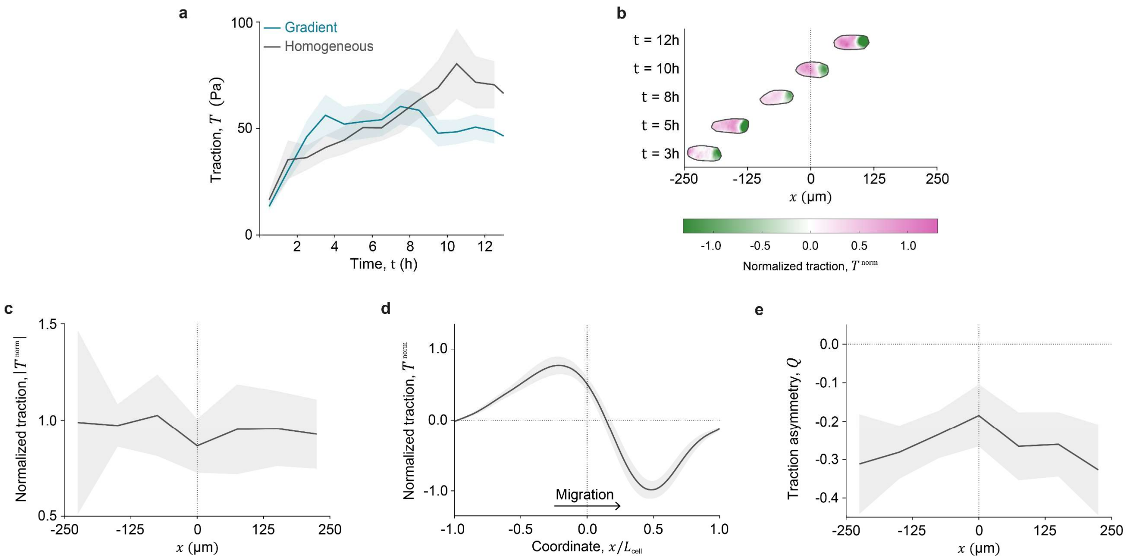


**Extended Data Figure 5 – Traction forces on homogeneous patterns. a,** Mean traction magnitude for cells migrating on homogeneous patterns (N=11) and gradient patterns (N=23). Error bars are SEM. **b,** Time lapse of the normalized traction map of a representative cell migrating on a homogeneous pattern. **c,** Mean normalized traction magnitude for cells migrating on homogeneous patterns (N=11). Error bars are SEM. **d,** Mean profiles of $x$ component of the normalized traction forces for cells migrating on homogeneous patterns (N=11). Error bars are SEM. **e,** Mean normalized traction quadrupole of cells migrating on homogeneous patterns (N=11). Error bars are SEM. Data were obtained from three independent experiments.

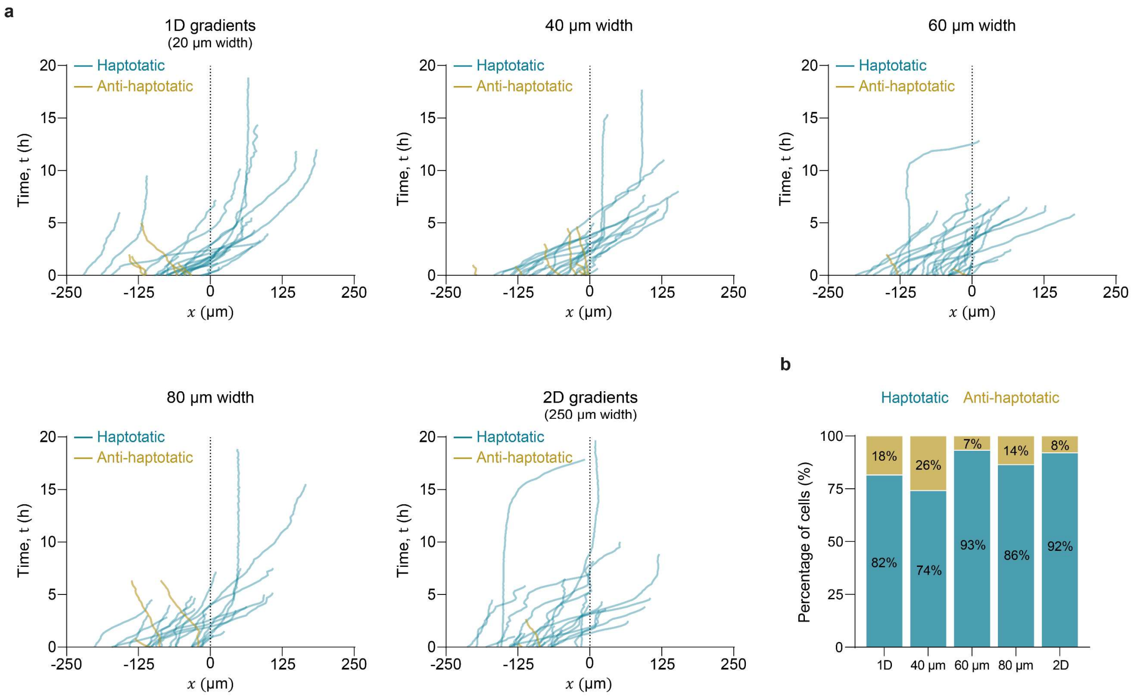


**Extended Data Figure 6 – Initial migration trajectories on gradients are independent of confinement width. a,** First run trajectories of cells migrating on gradients with widths of 20 μm (N=27), 40 μm (N=27), 60 μm (N=30), 80 μm (N=22) and 250 μm (N=25). **b,** Haptotaxis probability of cells migrating on gradients with widths of 20 μm (N=27), 40 μm (N=27), 60 μm (N=30), 80 μm (N=22) and 250 μm (N=25). 20, 40, 60 and 80 μm widths data were obtained from 6 replicates. 250 μm width data were obtained from 5 replicates.

#### Supplementary Theory Note

### 1 Differential friction model for haptotaxis

We first consider the dynamics of a single protrusion $i$ with a protrusive front at location $x_i(t)$ (Fig. 1). At the front, actin filaments polymerize against the cell membrane with a polymerization speed $v_p^{(i)}$. Actomyosin contraction within the cell creates an actin retrograde flow $v_r^{(i)}$ away from the cell edge. The cell membrane then advances with a speed $v_i$ determined by a balance of polymerization and retrograde flow:

$$v_i = v_p^{(i)} - v_r^{(i)} \tag{1}$$

Using force balance at the leading edge of the protrusion and the assumptions of the molecular clutch model, we will derive an expression for $v_r$ [1,2]. Specifically, backwards forces are given by the sum of contractile force $f_c$ (due to actomyosin contractility) and forces due to membrane tension $f_m$. These are locally balanced by a forward force arising due to the retrograde flow $f_f(v_r)$ [1] (Fig. 1):

$$f_f(v_r) = f_c + f_m \tag{2}$$

We will next make simplifying assumptions for each of these forces. First, according to the molecular clutch model, the force $f_f(v_r)$ exerted by the cell against the substrate is akin to a friction force exerted by adhesion molecules transiently bound to moving filaments. The kinetics of the number of adhesions $n_i$ are described by

$$\dot{n}_i = k_{on}(\rho(x_i))(N - n_i) - k_{off}n_i \tag{3}$$

where $k_{on}(\rho(x_i))$ is the binding rate of adhesions, $N$ the total number of binding sites and $k_{off}$ the unbinding rate. In general, this unbinding rate can be force-dependent (catch/slip bonds), which we neglect here. Instead, we focus on the dependence of the binding rate on the substrate ligand density $\rho(x)$. Assuming the adhesions act as linear elastic links, the backwards force acting on the substrate – and thus the forward force acting on the protrusion – is then

$$f_f(v_r) = \alpha v_r^{(i)} + \kappa n_i \langle \delta \rangle \tag{4}$$

where $\alpha$ is a bare friction coefficient, $\kappa$ is the stiffness of an adhesion and $\langle \delta \rangle$ is the average extension of an adhesion. Since adhesions remain bound for a typical time $1/k_{\text{off}}$, we can estimate their extension as

$$\langle \delta \rangle = \frac{v_r^{(i)}}{k_{\text{off}}} \tag{5}$$

Throughout, we will assume simplified adhesion dynamics. Specifically, we assume that the number of bound adhesions is in steady state and $N \gg n_i$, therefore $n_i =$

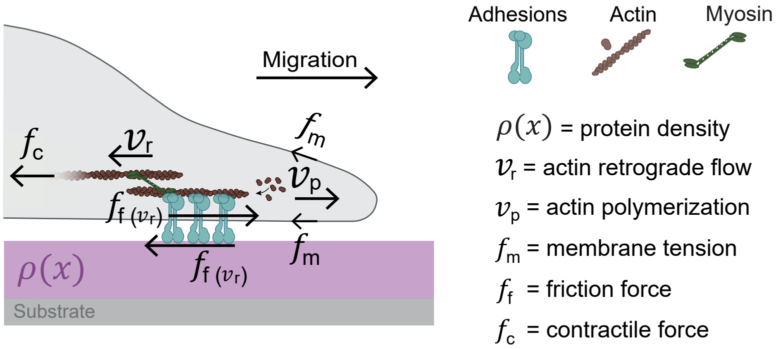


Figure 1: Schematic of a growing protrusion. Actin is polymerized with speed $v_p$. Contractile forces $f_c$ initiate retrograde actin flow $v_r$. Through coupling through adhesions, this flow exerts an effective frictional force $f_f(v_r)$ on the substrate, and an equal and opposite force onto the protrusion. The protrusion is coupled to the rest of the cell through a force due to membrane tension $f_m$.

$k_{on}(\rho)N/k_{\text{off}}$. Moreover, we assume that the binding rate of adhesions is proportional to the ligand density, $k_{on}(\rho) = k_{on}^{(0)}\rho$. Therefore,

$$f_f(v_r) = (\alpha + \beta\rho(x_i))v_r \tag{6}$$

with $\beta = \kappa k_{on}^{(0)} N/k_{\text{off}}^2$. For the contractile force $f_c$, we assume that it is proportional to the number of available actin filaments in the protrusion, which is determined by the balance of backward flow $v_r$ and depolymerization [3], and thus assume $f_c = \gamma v_r$.

For the force due to membrane tension $f_m$, we assume the simplest case of a global elastic cell coupling, $f_m = k(L - L_0)$ with equilibrium length $L_0$ and spring constant $k$.

Putting everything together and relabelling parameters $\alpha \to \alpha + \gamma$, we obtain the relation for retrograde flow:

$$v_r^{(i)} = \frac{k(L - L_0)}{\alpha + \beta\rho(x_i)} \tag{7}$$

Thus, we obtain the speed of a single protrusion as a function of local ligand density:

$$v_i = v_p^{(i)} - \frac{k(L - L_0)}{\alpha + \beta\rho(x_i)} \tag{8}$$

Our model therefore suggests that, at fixed polymerization speed, a protrusion at higher ligand density $\rho$ will exhibit slower retrograde flow, and thus will advance faster.

To obtain a closed dynamical system, we will additionally require a dynamical equation for $v_p^{(i)}$. We will derive this separately for two different scenarios in the next sections. First, as a model of the initial dynamics in the gradient, we will consider the dynamics of an initially unpolarized cell that has two protrusions left and right, whose competition gives rise to motion. Second, to model the long-term dynamics of cells and to generalize the model to 2D, we will consider a fully polarized cell with a single protrusion and non-zero average polarity.

#### 2 Initial spreading and migration of an unpolarized cell in a ligand density gradient

We first derive an equation of motion of an initially unpolarized cell in a non-uniform ligand density field $\rho(x)$, which grows to protrusions to the left and right. This allows us to study the behaviour of cells initially deposited on the gradient and their initial migration up/down the gradient.

Specifically, we consider two protrusions $i = \{1, 2\}$ with positions $x_2 < x_1$. We determine a dynamical equation for the speed of the center of mass of the cell $X = \frac{1}{2}(x_1 + x_2)$, given by $\dot{X} = \frac{1}{2}(v_1 - v_2)$ since we define $\dot{x}_2 = -v_2$ and $\dot{x}_1 = v_1$ to point in opposite directions.

We first derive the equation for cell motion:

$$\dot{X} = \tfrac{1}{2}(v_p^{(1)} - v_p^{(2)}) - \tfrac{1}{2}(v_r^{(1)} - v_r^{(2)}) \tag{9}$$

$$= \tfrac{1}{2}\Delta v_p - \tfrac{1}{2}\left(\frac{1}{\alpha + \beta\rho(x_1)} - \frac{1}{\alpha + \beta\rho(x_2)}\right) k(L - L_0) \tag{10}$$

To simplify the model, we linearize the gradient, i.e.

$$\rho(x_1) \approx \bar{\rho} + \tfrac{L}{2}\nabla\rho \tag{11}$$

$$\rho(x_2) \approx \bar{\rho} - \tfrac{L}{2}\nabla\rho \tag{12}$$

and assume that the gradient is small, i.e. $|\nabla\rho| \ll 2(\alpha + \beta\bar{\rho})/(\beta L)$, allowing an expansion of Eq. (10):

$$\dot{X} \approx \tfrac{1}{2}\Delta v_p + \frac{\beta k L(L - L_0)}{4[(\alpha + \beta\bar{\rho})]^3}\nabla\rho \tag{13}$$

Within the same approach, we can also derive the equation for the dynamics of length fluctuations:

$$\dot{L} = v_1 + v_2 \tag{14}$$

$$= v_p^{(1)} + v_p^{(2)} - (v_r^{(1)} + v_r^{(2)}) \tag{15}$$

$$\approx 2v_0 + \left(\frac{1}{\alpha + \beta\rho(x_1)} + \frac{1}{\alpha + \beta\rho(x_2)}\right) k(L - L_0) \tag{16}$$

$$\approx 2v_0 - \frac{2k(L - L_0)}{\alpha + \beta\bar{\rho}} \tag{17}$$

where in the first step, we neglected the fluctuations in $(v_p^{(1)} + v_p^{(2)})$ as the leading order contribution is the constant $2v_0$. In the second step, we use the small linear gradient assumption as above.

Based on our model, the friction with the substrate is highest in the region of highest ligand density, i.e. the highest local $\bar{\rho}$. This makes two predictions for experiment: first, at this location, the coupling of retrograde flow to the substrate is the most effective, meaning that the traction forces are highest. This is indeed observed experimentally (Fig. 2h). Since this also leads to the most productive outward protrusion growth, also the cell length is maximal in the region of highest average ligand density $\bar{\rho}$, as suggested by Eq. (17). This is also observed experimentally (Fig. 1e).

To obtain a closed dynamical system, we finally hypothesize that the polymerization velocity in each protrusion $i$ follows persistent dynamics with a typical speed $v_0$ around which it fluctuates with time-scale $\tau$:

$$\frac{dv_p^{(i)}}{dt} = -\frac{v_p^{(i)} - v_0}{\tau} + \frac{1}{\sqrt{2}}\xi(t) \tag{18}$$

where $\xi(t)$ is a Gaussian white noise with $\langle \xi(t) \rangle = 0$ and $\langle \xi(t)\xi(t') \rangle = 2D\delta(t - t')$. This results in persistent dynamics of the difference of polymerization speed:

$$\frac{d\Delta v_p}{dt} = -\frac{\Delta v_p}{\tau} + \xi(t) \tag{19}$$

Taking the difference in polymerization speeds $\Delta v_p$ to be indicative of the cell polarity $P$, we relabel $P = \frac{1}{2}\Delta v_p$. For simplicity, we have approximated cell length to remain constant and absorbed cell length variation into the gradient coupling parameter $\chi = \beta k L(L - L_0)/4[(\alpha + \beta\bar{\rho})]^3$. This yields the equations of motion for the migration of the cell in a gradient of ligand density

$$\dot{X} = P + \chi \nabla \rho \tag{20}$$

$$\dot{P} = -\frac{P}{\tau} + \xi(t) \tag{21}$$

Here the second term in the first equation is the coupling of cell dynamics to the density gradient. This means that the clutch model predicts haptotaxis in the following way: on larger average density, the speed of the a growing protrusion is higher and therefore the cell velocity reorients to align with the gradient of density, i.e. moving up density gradients. Mathematically, this term is akin to the drift term in the Keller-Segel model of chemotaxis.

Interestingly, this model suggests a number of non-trivial correlations emerging from persistence, i.e. from the fact that cells that reached a specific position are there because of a specific polarity history. For instance, although the model only couples actin retrograde speed $v_r$ to local fibronectin density $\rho$ (as observed on homogeneous substrates), simulations also show correlations between $P$ (i.e., $\Delta v_p$) and gradients of $\rho$, as seen experimentally (Fig. 2e).

#### 3 Numerical implementation and parameter overview

We numerically integrate the equations of motion using custom-written python code using numpy [4]. We use a stochastic Euler scheme with small time-step $\delta t$:

$$X(t + \delta t) = X(t) + (P(t) + \chi \nabla \rho(X(t)))\delta t \tag{22}$$

$$P(t + \delta t) = P(t) + [-P(t)/\tau]\delta t + \sqrt{2D} r_t \sqrt{\delta t} \tag{23}$$

where $r_t \sim \mathcal{N}(0, 1)$ is a zero-mean, unit-variance normally distributed random numbers. We use a time interval $\delta t = 0.005$h, iterated for $N_t = 4000$ time steps, equating to a total trajectory length of 20h like in the experiment.

The model has 3 parameters: $\tau$, $D$ and $\chi$. We first use the experiments of cells migrating on uniform density substrates (on which $\chi$ is irrelevant since $\nabla \rho = 0$), to fit the parameters $\tau$ and $D$. Using these values, we then vary the magnitude of $\chi$ to match the proportion of cells initially migrating up/down the gradient (Fig. 2). Since $\rho(x)$ is measured in ng/cm$^2$, which we convert to ng/$\mu$m$^2$, the units of $\chi$ are $\mu$m$^4$ ng$^{-1}$ h$^{-1}$.

In the main text, we then use the model to make a number of additional predictions on the behaviour of the trajectories beyond the initial directionality up/down the gradient. The parameters used throughout the figures are specified in Table 1.

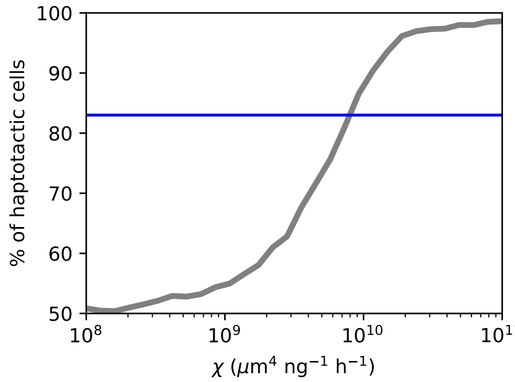


Figure 2: Proportion of cells initially migrating up the gradient as a function of $\chi$. The experimentally measured proportion of $\approx$ 83% is indicated as a blue horizontal line.

| **Symbol** | **Value** | **Units** |
|---|---|---|
| $D$ | 800 | $\mu$m$^2$ h$^{-3}$ |
| $\tau$ | 2 | h |
| $\chi$ | $7 \times 10^9$ | $\mu$m$^4$ ng$^{-1}$ h$^{-1}$ |

Table 1: Overview of parameters used in the 1D model.

Equivalent parameters are used in the 2D model.

#### 4 Dynamics of a polarized cell in a ligand density gradient

We finally extend the model to the dynamics of polarized cells that exhibit a single leading edge at the front $f$, and a passive back $b$. Thus, we assume that $v_p^{(b)} \approx 0$, while $v_p^{(f)}$ follows dynamics with a non-zero steady-state polarity

$$\frac{dv_p^{(f)}}{dt} = v_p^{(f)} - \frac{(v_p^{(f)})^3}{v_0^2} + \sigma\xi(t) \tag{24}$$

To implement our model in 2D, we straightforwardly generalize Eq. (20):

$$\dot{\mathbf{X}} = \mathbf{P} + \chi \boldsymbol{\nabla} \rho \tag{25}$$

For cells with a non-zero steady-state polarity, for computational simplicity, we assume a constant polarity vector $\mathbf{P} = v_0(\cos\theta, \sin\theta)$ with fluctuating orientation $\theta$. In our 2D model, we additionally have to capture the cell confinement. Since cells in narrow confinements (such as the 20-40 $\mu$m lines) tend align their long axis, and thus their polarity, with the long axis of the system (though not necessarily with the gradient), we include an alignment term with width-dependent amplitude in the dynamics of the polarity angle $\theta$:

$$\dot{\theta} = C(w)\sin 2\theta + \xi(t) \tag{26}$$

where $\langle \xi(t) \rangle = 0$ and $\langle \xi(t)\xi(t') \rangle = 2D_r\delta(t-t')$ with $D_r$ the rotational diffusion coefficient. As a phenomenological description of the decreasing strength of lateral confinement with width, we take $C(w) = C_0 e^{w/w_0}$ with $C_0 = 3$ and $w_0 = 40$ $\mu$m.
